# Characterization of γ-Glutamyl Peptidases and γ-Glutamyl Cyclotransferases for Glutathione Degradation in *Arabidopsis*

**DOI:** 10.1101/2021.10.29.466391

**Authors:** Takehiro Ito, Taisuke Kitaiwa, Kosuke Nishizono, Minori Umahashi, Shunsuke Miyaji, Shin-ichiro Agake, Kana Kuwahara, Tadashi Yokoyama, Akiko Maruyama-Nakashita, Ryosuke Sugiyama, Masami Yokota Hirai, Naoko Ohkama-Ohtsu

## Abstract

- Organic sulfur is stored as glutathione (GSH) in plants. In *Arabidopsis*, γ-glutamyl cyclotransferases (GGCT2;1, GGCT2;2, and GGCT2;3) degrade cytosolic GSH, but they do not fully explain the rapid GSH turnover. Here, we demonstrate that γ-glutamyl peptidases, GGP1 and GGP3, play a substantial role in degrading GSH in the cytosol.
- We conducted yeast complementation assay and activity assay of recombinant proteins to identify the novel GSH degradation enzymes. The expression patterns were investigated by RT-qPCR. GSH concentrations in the mutants were also analyzed.
- *GGP1* complemented the yeast phenotype. Recombinant GGP1 and GGP3 showed reasonable *K_m_* values considering cytosolic GSH concentration, and their activity was comparable to that of GGCTs. The *GGP1* transcript was highly abundant in mature organs such as rosette leaves. The expression of *GGCT2*;*1* was conspicuously enhanced under sulfur deficiency. GSH concentration was higher in *ggp1* knockout mutants regardless of nutritional conditions; the concentration was higher in *ggct2*;*1* knockout mutants under sulfur-deficient conditions.
- We propose a model wherein cytosolic GSH is degraded fundamentally by GGP1. The degradation is accelerated by GGCT2;1 under sulfur deficiency. Given the energy cost throughout the reactions, GGPs could render a more efficient route for GSH degradation than GGCTs.

## Introduction

Glutathione (GSH; L-γ-glutamyl-L-cysteinyl-glycine) plays a variety of essential roles in plants. GSH reversibly undergoes redox reactions through its thiol residue and maintains the redox status of cells. When oxidized, two glutathione molecules form dimer and become oxidized glutathione (GSSG), which is returned to the reduced GSH by glutathione reductase. GSH is essential in alleviating oxidative stress in plants since most GSH in the cell is in its reduced form and can reduce oxidants (Foyer and Noctor, 2011; Noctor *et al*., 2011). GSH is also involved in the redox regulation of proteins (Gao *et al*., 2009). Phytochelatins, the polymerized forms of GSH, contribute to plant tolerance toward heavy metals and metalloids, such as cadmium and arsenate (Cobbet, 2000; Mendoza-Cózatl, 2011). GSH also plays another role in the storage of reduced sulfur in cells. Sulfur is taken up by plant cells mainly as sulfate, which is then reduced to sulfide. Sulfide can be incorporated into the amino acid skeleton of *O*-acetyl-L-Ser to produce Cys, which is the donor for all further metabolites (Koprivova and Kopriva, 2014). Cys, however, is not a storage form of reduced sulfur because extra concentration of cysteine is toxic due to its high reactivity (Romero *et al*., 2014; Deshpande *et al*., 2017). Cys is incorporated into GSH, which exists in high concentrations in plant tissues, acts as a substrate for central biochemicals, and maintains Cys availability for protein synthesis. Thus, GSH acts as a precursor for metabolites that, in turn, may be related to the GSH degradation rate (Leustek *et al*., 2000). GSH concentrations in the phloem sap are in the mM range, and GSH is considered the transport form of reduced sulfur (Kuzuhara *et al*., 2000).

GSH is synthesized from its constituent amino acids via a two-step ATP-dependent reaction. First, γ-glutamylcysteine (γ-EC) is synthesized from Glu and Cys by γ-EC synthetase (Hell and Bergmann, 1990), which is encoded by *GSH1* (May and Leaver, 1994), and Gly is incorporated by GSH synthetase encoded by *GSH2* (Wang and Oliver, 1996). However, GSH degradation pathways are not as straightforward as this synthesis pathway. In mammals, GSH is degraded during the γ-glutamyl cycle (Supporting Information Fig. **S1**; Orlowsk and Meister, 1970; Meister and Larsson, 1995). In this cycle, GSH is degraded outside cells by an exoenzyme, γ-glutamyl transpeptidase (GGT), which hydrolyzes GSH and transfers the Glu residue to water or other amino acids. When transferred to water, Glu is produced; when transferred to amino acids, γ-glutamyl amino acids are produced. Glu and γ-glutamyl amino acids are incorporated into cells, and γ-glutamyl amino acids are processed by cytosolic enzyme, γ-glutamyl cyclotransferase (GGCT). This enzyme detaches the γ-glutamyl residue of γ-glutamyl amino acids, and releases it as 5-oxoproline. 5-oxoproline can be converted to Glu by another cytosolic enzyme, oxoprolinase. The other reaction product of GSH degradation by GGT is cysteinylglycine (Cys-Gly). This dipeptide is degraded into Cys and Gly by dipeptidases. In this way, GSH is degraded into three constituent amino acids, Glu, Cys, and Gly, and they are utilized for biosynthesis of proteins or GSH in the cells (Fig. S1).

In *Arabidopsis*, the turnover of GSH is as high as 80% in 1 d under normal conditions (Ohkama-Ohtsu *et al*., 2008), and genes responsible for the degradation pathway have been extensively explored. There are four homologs of the mammalian GGTs: GGT1, 2, 3, and 4. GGT1 and GGT2 are localized to the apoplast and GGT4 to the vacuole (Grzam *et al*., 2007; Martin *et al*., 2007; Ohkama-Ohtsu *et al*., 2007a, b). Among them, GGT1 and GGT4 are actively expressed in the whole plant body, so a *ggt1/ggt4* knockout mutant, which shows no GGT activity, was analyzed in detail (Ohkama-Ohtsu *et al*., 2008). However, GSH degradation rate and GSH concentration were not affected by the defect in GGT activity. This strongly suggested that GGT is not involved in GSH degradation; instead, GGP1 is responsible for GSSG degradation (Ohkama-Ohtsu et al., 2007a) and GGT4 is for GSH-conjugate degradation (Grzam et al., 2007; Ohkama-Ohtsu et al., 2007b). After all, other GSH degradation pathways needed to be identified.

Ohkama-Ohtsu *et al*. (2008) also found that 5-oxoproline levels are correlated with GSH or γ-EC levels, and proposed that GGCT, a cytosolic enzyme which degrades γ-glutamyl amino acids and releases 5-oxoproline, also degrades GSH. In addition, activity assay using protein extract suggested that 5-oxoproline is more likely to be supplied by GSH degradadtion than by γ-EC degradation (Ohkama-Ohtsu *et al*., 2008). Later, GGCT which has GSH degradation activity was indeed identified in mammals (Kumar *et al*., 2012), and three homologs were identified in *Arabidopsis*: *GGCT2*;*1*, *GGCT2*;*2*, and *GGCT2*;*3*. All of them degrade physiological concentration of GSH into 5-oxoproline and Cys-Gly *in vitro* and complement the GSH degradation-defective yeast mutant (Paulose *et al*., 2013; Kumar *et al*., 2015). Among these *GGCT* genes, expression patterns of *GGCT2*;*1* have been analyzed in detail. *GGCT 2*;*1* mRNA is rapidly accumulated under various conditions: pollen tube growth (Want *et al*., 2008), heavy metal stress (Kovalchuk *et al*., 2005), salinity stress (Gong *et al*., 2005), and sulfur starvation (Maruyama-Nakashita *et al*., 2005; Hubberten *et al*., 2012; Bielecka *et al*., 2014). Paulose *et al*. (2013) reported that the promoted *GGCT2*;*1* expression under arsenite stress can contributes to heavy metal tolerance by recycling Glu in GSH and saving the energy for *de novo* Glu synthesis. Joshi *et al*. (2019) also reported that GGCT2;1 is involved in the regulation of root architecture under sulfur deficiency by recycling Cys in GSH. However, what is responsible for the rapid turnover of GSH is still unclear because the expression of *GGCT2*;*1* is very low under normal conditions. We hypothesized that enzymes other than GGCT2;1 contribute to cytosolic GSH degradation under normal conditions.

In the present study, we show that the cytosolic enzymes γ-glutamyl peptidase 1 (GGP1) and GGP3 have GSH degradation activity comparable to that of GGCT2;1, GGCT2;2, and GGCT2;3. Because GGP1 and GGP3 also process GSH-conjugates in the biosynthesis of glucosinolates and camalexin (Geu-Flores *et al*., 2009), they are considered bifunctional enzymes involved in GSH degradation and glucosinolates and camalexin synthesis. We further suggest that cytosolic GSH is fundamentally degraded by GGP1, except that GGCT2;2 is crucial in developing organs and GGCT2;1 is vital under sulfur deficiency.

## Materials and Methods

### Yeast strains and screening

The *Saccharomyces cerevisiae* strains used in this study, listed in Table **S1**, were obtained from EUROSCARF (http://www.euroscarf.de/). Strains Y05729 (*dug2Δ*) and Y05729 (*dug3Δ*) were constructed from BY4741 by insertional mutagenesis (Winzeler *et al*., 1999). All strains were missing *MET15* and were unable to assimilate inorganic sulfur.

An *A. thaliana* cDNA library was cloned into the expression plasmid pFL61 and transformed into competent yeast cells of *dug2Δ* or *dug3Δ* strains (Minet *et al*., 1992) using the lithium acetate method (Ito *et al*., 1983). The transformants were screened on minimal selection plates (SD-ura), in which all sulfate ions were replaced with chloride ions and supplemented with 200 μM GSH. All plates were incubated at 28 °C for 3–5 d. The transformation efficiency was calculated from yeast grown on plates (SD-ura) supplemented with 200 μM methionine. The plasmids transformed into yeast were extracted using Zymoprep™ Yeast Plasmid Miniprep I (Zymoresearch, http://www.zymoresearch.com/).

The inserts in the clones were sequenced with the ABI PRISM 3130 and 3500 Genetic Analyzer (Applied Biosystems), and primers pGK5 and pGK3 listed in Table **S2** (Nozawa *et al*., 2006).

### Construction of the plasmids for transformation into yeast

Total RNA was extracted from *A. thaliana* using an RNeasy Plant Mini Kit (Qiagen, Hilden, Germany) followed by cDNA synthesis with SuperScript III reverse transcriptase (Invitrogen, https://www.thermofisher.com/) and the Oligo (dT)_16_ primer. The ORF sequences for *GGP1* (*AT4G30530*), *GGCT2*;*1* (*AT5G26220*), *GGCT2*;*2* (*AT4G31290*), and *GGCT2*;*3* (*AT1G44790*) were amplified from the *A. thaliana* cDNA template using the primer sets listed in Table **S2**. The amplified product for *GGP1* was subcloned into the *EcoRI-XhoI* site, and those for *GGCT2*;*1*, *GGCT2*;*2*, and *GGCT2*;*3* were subcloned into the *BamHI-EcoRI* site of the p416TEF yeast expression vector (Mumberg *et al*., 1995). These plasmids were transformed into *dug2Δ* and *dug3Δ* yeast strains using the lithium acetate method (Ito *et al*., 1983).

### Construction of *E. coli* expression strains

The coding sequences (CDSs) of *GGP1*, *GGP3*(*AT4G30550*), *GGCT2*;*1*, *GGCT2*;*2* and *GGCT2*;*3* were obtained from the *Arabidopsis* Information Resource (https://www.arabidopsis.org/), and they were amplified by PCR using KOD-Plus-Neo (TOYOBO, Osaka, Japan) with the primers listed in Table **S3**. The PCR products were cloned into the entry vector pENTR/D-TOPO (Invitrogen, MA, USA), and the sequences of inserts were confirmed by Eurofins Genomics (Tokyo, Japan) with M13 primers (Table **S3**). Subsequently, each CDS was cloned into the destination vector pDEST17 (Invitrogen, MA, USA) via the LR recombination reaction. Finally, the constructed plasmids were transformed into *E. coli* expression strain BL21-AI (Invitrogen, MA, USA). All operations were performed according to the manufacturers’ protocols.

### Expression and purification of the recombinant proteins

BL21-AI possessing each construct was cultured in LB medium containing 100 mg/ml ampicillin up to O.D._600_ ~ 0.6 at 37 °C at 150 rpm. Then, L-arabinose was added to a final concentration of 0.02% (w/v), and the cells were cultured for another 17 h at 18 °C at 120 rpm. Cells were harvested by centrifugation at 2500 × *g* for 5 min at 4 °C. The target proteins were obtained as N-terminal His-tagged proteins, followed by purification via His60 Ni Gravity Column Purification Kit (Clontech Laboratories, CA, USA), according to the manufacturer’s instructions; except that 1 mM phenylmethanesulfonyl fluoride (PMSF) was added to the extraction and wash buffers of the kit. Out of the five proteins expressed in *E. coli*, GGP1, GGP3, and GGCT2;3 were collected from the soluble fraction and utilized for the subsequent experiments. The proteins were dialyzed overnight in a buffer containing 50 mM Tris-HCl (pH 8.0) and 150 mM NaCl at 4 °C. The results were confirmed using SDS-PAGE (Fig. **S2**). Protein concentrations were determined using the Bradford reagent (Bio-Rad, CA, USA) and bovine serum albumin as the standard.

### Activity assay of recombinant proteins for GSH or other γ-glutamyl compounds

Kinetic parameters were determined using 0.5 μg of GGP1, 0.25 μg of GGP3, or 2.2 μg of GGCT2;3. The proteins were incubated in 50 μl of reaction mixture containing 50 mM Tris-HCl (pH 8.0) and 0.25 mM to 15.0 mM GSH for 30 min at 37 °C, and the reaction was terminated by adding 10 μl of 1.5 M HCl. The solution was centrifuged at 11,000 × *g* for 15 min at 4 °C, and the Cys-Gly released in the supernatant was quantified by HPLC as described below. Kinetic parameters were calculated using GraphPad Prism 9 software (GraphPad Software, CA, USA). For degradation activity for other γ-glutamyl compounds, 0.5 μg of GGP1 or 0.25 μg of GGP3 was incubated in 50 μl of reaction mixture containing 50 mM Tris-HCl (pH 8.0) and 5.0 mM GSH, 5.0 mM γ-Glu-Cys (Sigma-Aldrich), 5.0 mM □-Glu-Ala (Sigma-Aldrich), or 2.5 mM GSSG. The reaction was conducted as described previously. As described below, released Cys-Gly was quantified for GSH, Cys was quantified for γ-Glu-Cys, and Ala was quantified for γ-Glu-Ala.

### Analysis of Ala

Ala in the reaction solutions of activity assays was derivatized with the AccQ-Fluor Reagent Kit (Waters, http://www.waters.com/waters/home) and measured by HPLC using an AccQ-Tag column (Waters) according to the manufacturer’s instructions.

### Thiol analysis with HPLC

Thiols extracted from plant tissues with 0.01 N HCl, or in the reaction solutions of activity assays, were quantified by HPLC after derivatization with monobromobimane (Minocha *et al*., 2008; Nishida *et al*. 2016). The HPLC system (Shimadzu, Japan) consisted of a system controller (CBM-20A), two pumps (LC-20AD), a degasser (DGU-20A_3_), an autosampler (SIL-20A), a column oven (CTO-20AC), and a fluorescence detector (RF-20A). The column was a Shim-pack FC-ODS (4.6 mm × 150 mm; Shimadzu, Japan).

### Hydroponic culture of plants for sampling at vegetative and reproductive stages

*Arabidopsis thaliana* Col-0 wild-type plants were grown hydroponically in 50 ml conical tubes at 22 °C under 16 h light (100 μmol^-1^m ^-2^ s^-1^) / 8 h dark as described by Ohkama-Ohtsu *et al*. (2007b) using MGRL medium (Fujiwara *et al*., 1992; Hirai *et al*., 1995). The vegetative-stage samples were grown for 22 d, three oldest leaves from the first true leaf were collected as mature leaves, and three youngest leaves were collected as young leaves. For the reproductive stage, samples were grown for 33 d, and flower + buds, siliques 1–2 cm in length, stems, and all cauline and rosette leaves were collected separately. Roots were divided into three equally long parts, and the upper, middle and lower roots were collected separately. Samples were immediately frozen with liquid nitrogen and stored at −80 °C until RNA extraction.

### Liquid culture of plants in flasks

*Arabidopsis* plants were grown in flasks, as described in Xiang and Oliver (1998) using MGRL medium (Fujiwara *et al*., 1992). For sulfur-deficient treatments, all sulfate ions in the MGRL medium were replaced with an equal molar concentration of chloride ions (Hirai *et al*., 1995). In nitrogen-deficient treatments, all the nitrate ions were replaced with an equal molar concentration of chloride ions. In the nitrogen-deficient experiment, 12 d-old plants grown in a standard medium were transferred to a nitrogen-deficient medium and grown for an additional 2 d. In sulfur-deficient experiments, 10 d-old plants grown in the standard medium were transferred to sulfur-deficient medium and grown for an additional 4 d. Control plants were grown for 14 d in a standard MGRL medium, and the medium was replaced after 10 d. The entire bodies of plants in each flask were pooled immediately frozen with liquid nitrogen and stored at −80 °C until analysis.

### RNA extraction and quantitative PCR analysis

RNA was extracted from frozen samples using the RNeasy Plant Mini Kit (Qiagen, Germany), followed by reverse transcription using the PrimeScript™ RT Reagent Kit with gDNA Eraser (Perfect Real Time) (Takara Bio, Shiga, Japan). Quantitative PCR was conducted using the KAPA SYBR FAST qPCR Master Mix (2X) Kit (KAPA Biosystems, MA, USA) and LightCycler 96 (Roche Diagnostics, Switzerland) following the manufacturer’s protocols. The absolute quantification method was adopted using serial dilutions of DNA with known copy numbers for standard curves. The copy number of *β-tubulin* (*TUB4*, AT5G44340) was determined as the internal standard. The primers used are listed in Table **S4**. The normalized copy number was determined by dividing copy number of each gene by that of *β-tubulin*.

### Screening of *Arabidopsis* T-DNA knockout mutants for *ggp1* and *ggct2*;*1*

Unless otherwise indicated, *Arabidopsis thaliana* plants were grown on rockwool in a plastic container at 22 °C under a 24 h photoperiod (150 μmol m^-2^ s^-1^) under 16 h light/8 h dark conditions to establish mutant lines honoring T-DNA homozygously. The primers used for screening *ggp1* and *ggct2*; *1* mutant plants are shown in Table **S5**.

The *ggp1-1* mutant, which was used by Geu-Flores *et al*. (2011), was provided by GABI-kat (http://www.gabi-kat.de/, stock no. GK-319F10) (Rosso *et al*., 2003). Homozygous plants were screened by PCR using the gene-specific primers GK-319-F, GK-319-R and the T-DNA left-border primer GABI-8474 (http://www.gabi-kat.de/). A lack of amplifiable transcripts of *At4g30530* in the *ggp1-1* mutant was reported by Geu-Flores *et al*. (2011), and we also confirmed by quantitative RT-PCR using primers At4g30530F and At4g30530R. The *ggp1-2* mutant (SALK_089634) was provided by the Salk Institute (Alonso *et al*., 2003) and was obtained from the *Arabidopsis* Biological Resource Center (ABRC, https://abrc.osu.edu/). Homozygous plants were screened using the gene-specific primers SALK_089634F, SALK_089634R, and the T-DNA left-border primer pROKr3 (Lin and Oliver, 2008). The lack of an amplifiable transcript of *At4g30530* in the *ggp1-1* mutant was confirmed by quantitative RT-PCR using primers At4g30530F and At4g30530R.

The *ggct2*;*1-2* mutant (SALK_56007), which was used by Joshi *et al*. (2019), was provided by the Salk Institute (Alonso *et al*., 2003) and was obtained from the Arabidopsis Biological Resource Center (ABRC, https://abrc.osu.edu/). Homozygous plants were screened by PCR using the gene-specific primers GGCT2,1F, GGCT2,1R, and the T-DNA left-border primer pROKr3 (Lin and Oliver, 2008). A lack of an amplifiable transcript of *At5g26220* in the *ggct2*;*1-2* mutant was reported by Joshi *et al*. (2019), and we also confirmed by RT-PCR using the primers At5g26220F and At5g26220R.

## Results

### Isolation of *A. thaliana* cDNA clones that complement yeast *dug2Δ* or *dug3Δ* mutants

An *A. thaliana* cDNA library built in the yeast expression vector pFL61 (Minet *et al*., 1992) was transformed into the *Saccharomyces cerevisiae* yeast mutants *dug2Δ* or *dug3Δ* (DUG: Defective in Utilization of GSH; Ganguli *et al*., 2007). In *Saccharomyces cerevisiae*, three mutants, *dug1Δ*, *dug2Δ*, and *dug3Δ*, are known to be defective in GSH utilization (Ganguli *et al*, 2007). (DUG2-DUG3)_2_ complex first degrades GSH into Glu and Cys-Gly, and then the (DUG1)_2_ homodimer degrades Cys-Gly into Cys and Gly (Kaur *et al*., 2009, 2012). As a result of this transformation, *c*. 4.6 × 10^4^ yeast transformants were obtained in *dug2Δ*, and two independent colonies were isolated from the minimal selection plates with GSH as the sole sulfur source. From the *c*. 6.9 × 10^4^ yeast transformants in *dug3Δ*, three independent colonies were isolated under the same growth conditions. Plasmids were isolated from these colonies, and cDNAs expressed from the library were sequenced. All clones that complemented the *dug2Δ* and *dug3Δ* yeast mutants corresponded to the gene *γ-glutamyl peptidase 1 (GGP1*), which was initially identified as an enzyme processing GSH-conjugates in the biosynthesis of glucosinolates and camalexin (Geu-Flores *et al*., 2009).

To confirm the complementation of *dug2Δ* and *dug3Δ* yeast with *GGP1*, the open reading frame (ORF) of *GGP1* was cloned into the yeast expression vector p416TEF (Mumberg *et al*., 1995) and transformed into *dug2Δ* and *dug3Δ* mutants. Both yeast mutants transformed with the *GGP1* ORF grew on the medium when GSH was the sole sulfur source (Fig. **1**). This result suggested that GGP1 from *Arabidopsis* is sufficient to degrade GSH, whereas yeast DUG2 and DUG3 need to form an active complex. The genes *GGCT2*;*1*, *GGCT2*;*2*, and *GGCT2*;*3* were also cloned into p416TEF and were introduced into *dug2Δ* or *dug3Δ* mutants. As previously demonstrated, all GGCT genes complemented both yeast mutants (Fig. **1**) (Kumar *et al*., 2015).

**Fig. 1.**
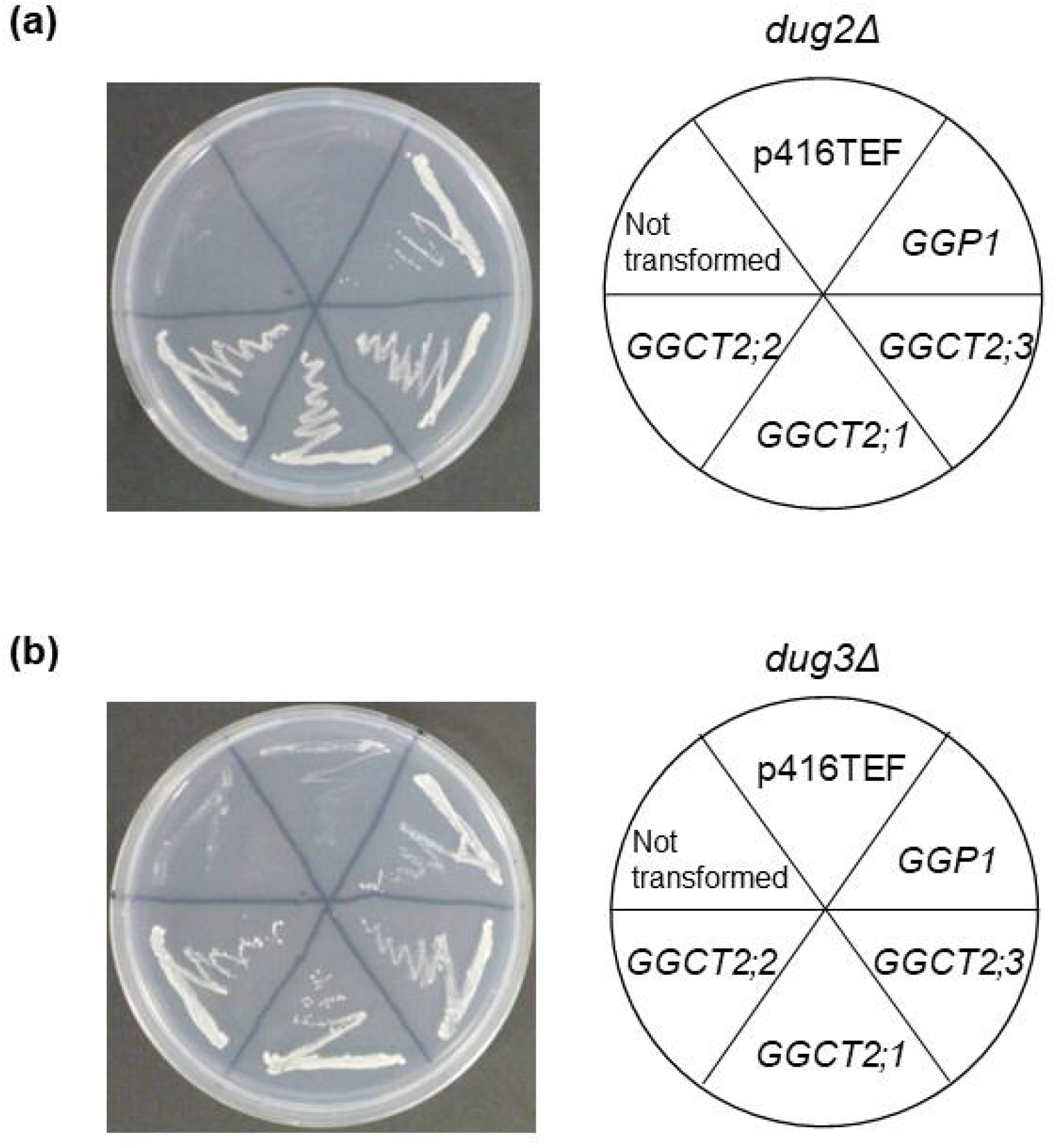
Functional complementation assay of *Arabidopsis γ-glutamyl peptidase 1* (*GGP1*), *γ-glutamyl cyclotransferase 2*;*1* (*GGCT2*;*1*), *GGCT2*;*2* and *GGCT2*;*3* in yeast mutants. The *Saccharomyces cerevisiae dug2Δ* (a) or *dug3Δ* (b) mutants (DUG: Defective in Utilization of Glutathione) were transformed with plasmids carrying the *Arabidopsis* genes *GGP1*, *GGCT2*;*1*, *GGCT2*;*2*, or *GGCT2*;*3* driven by the TEF promoter in the p416TEF vector. Yeast transformants were grown in an SD medium supplemented with glutathione (GSH). The vector p416TEF was used as a control, and the growth of untransformed *dug2Δ* (a) or *dug3Δ* (b) is also shown.

### Degradation activities of recombinant proteins from *E. coli*

We investigated the enzymatic properties of GGPs and GGCTs using recombinant proteins. There are five *GGP* genes in *Arabidopsis*, but the expression of *GGP2*, *GGP4*, and *GGP5* is very low, and only *GGP1* and *GGP3* are actively expressed across all tissues (Geu-Flores *et al*., 2011); therefore, in addition to GGP1, GGP3 was analyzed in this experiment.

GGP1, GGP3, GGCT2;1, GGCT2;2, and GGCT2;3 were expressed in *E. coli*. GGP1, GGP3, and GGCT2;3 collected from the soluble fraction were utilized for the subsequent measurement of enzyme kinetis, and anlyzed together with the reported properties of GGCT2;1 and GGCT2;2 (Paulose *et al*., 2013; Kumar *et al*., 2015). The activity and affinity of the proteins were analyzed with various GSH concentrations ranging from 0.25 mM to 15.0 mM (Fig. **S3**; Table **1**). The *K_m_* of GGP1, GGP3, and GGCT2;3 was 5.0 mM (GGP1), 3.7 mM (GGP3), and 6.7 mM (GGCT2;3), respectively; their *k_cat_* was 19.9 min^-1^ (GGP1), 70.3 min^-1^ (GGP3), and 18.5 min^-1^ (GGCT2;3), respectively.

**Table 1.**
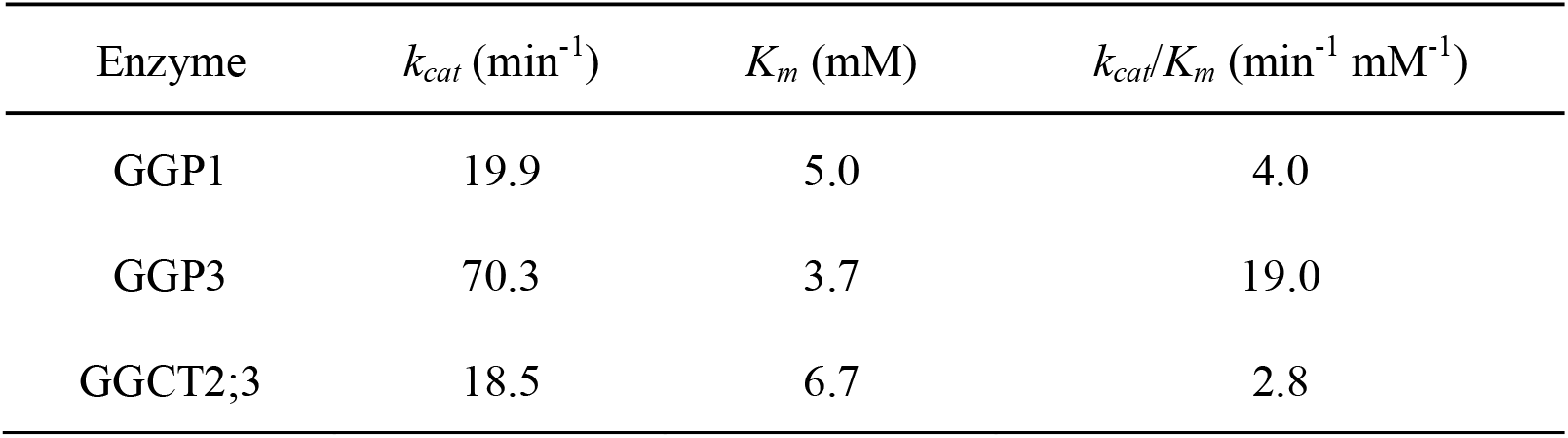
Kinetic parameters of GGP1, GGP3, and GGCT2;3.

| Enzyme | $k_{cat}$ (min<sup>-1</sup>) | $K_m$ (mM) | $k_{cat}/K_m$ (min<sup>-1</sup> mM<sup>-1</sup>) |
| --- | --- | --- | --- |
| GGP1 | 19.9 | 5.0 | 4.0 |
| GGP3 | 70.3 | 3.7 | 19.0 |
| GGCT2;3 | 18.5 | 6.7 | 2.8 |

In addition to that toward GSH, degradation activity of these enzymes toward other γ-glutamyl compounds—namely, oxidized glutathione (GSSG), γ-glutamyl-cysteine (γ-Glu-Cys), and γ-glutamyl-alanine (γ-Glu-Ala)—was also analyzed (Fig. **S4**). However, the activity of GGCT2;3 toward γ-Glu-Cys and γ-Glu-Ala was excluded from the analysis because it is known to be much lower than that toward GSH (Kumar *et al*., 2015). The results showed that GGP1, GGP3, and GGCT2;3 hardly degraded γ-Glu-Cys, γ-Glu-Ala, or GSSG.

### Comparison of expression levels of *GGP* and *GGCT* genes among organs

The distribution of *GGP1*, *GGP3*, *GGCT2*;*1*, *GGCT2*;*2*, and *GGCT2*;*3* was clarified by dividing plants at the vegetative and reproductive stage into 10 parts and analyzing the expression levels of these genes by quantitative RT-PCR using the absolute quantification method. The components of the 10 parts were the mature and young leaves of vegetative-stage plants and the flower/buds, siliques, stems, cauline and rosette leaves, and the upper, middle, and lower roots of reproductive-stage plants. In general, all *GGP* and *GGCT* genes, except *GGCT2*;*1*, were constitutively expressed in all organs (Fig. **2**). In particular, *GGP1* transcripts in rosette leaves were abundant. In contrast, the expression of *GGCT2*;*1* was much lower than that of other four genes in all organs.

**Fig. 2.**
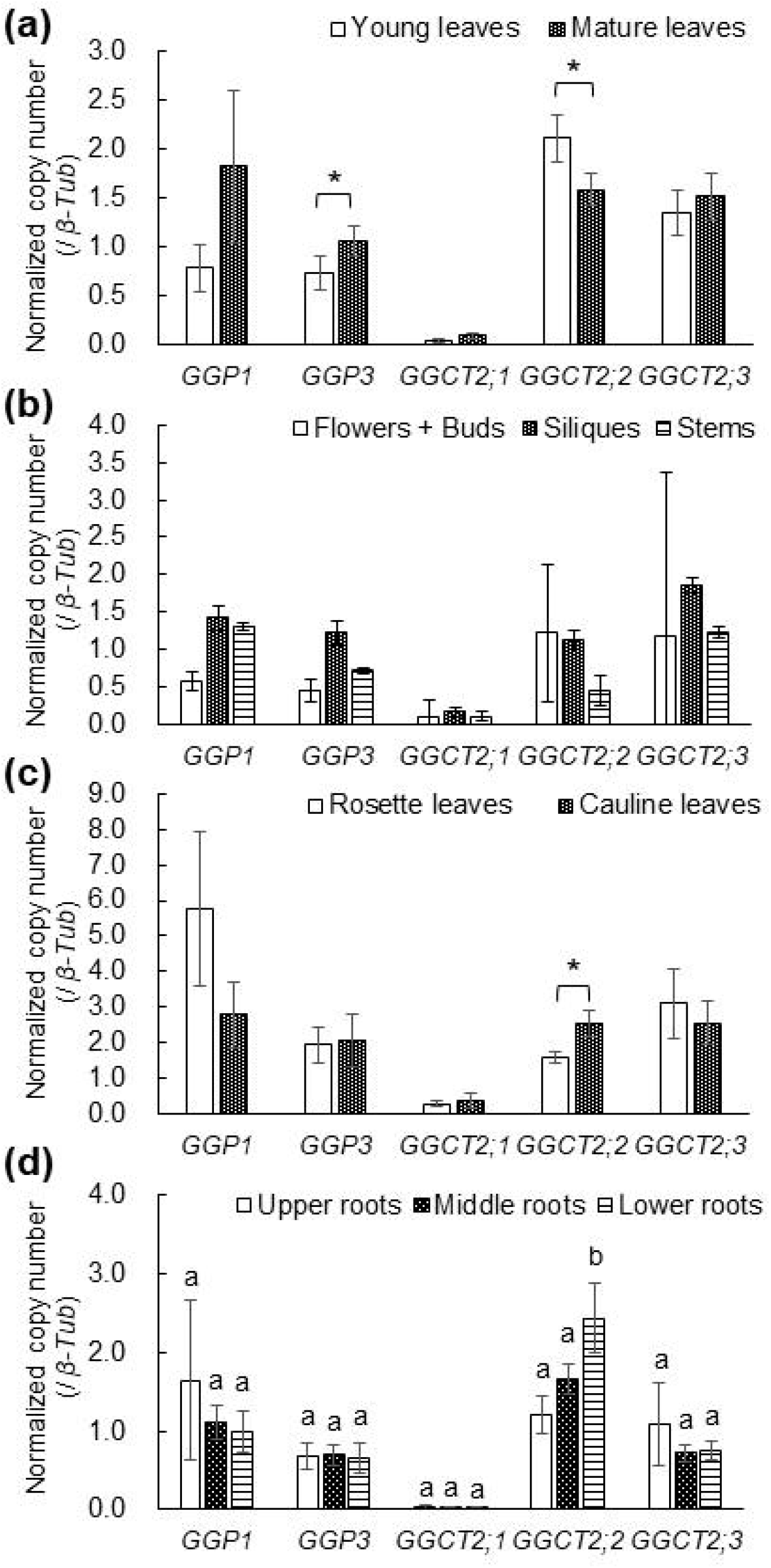
Normalized copy numbers of *GGP1*, *GGP3*, *GGCT2*;*1*, *GGCT2*;*2*, and *GGCT2*;*3* transcripts in various organs of wild-type plants. Plants were grown in hydroponic culture. Transcript copy numbers of *GGP1*, *GGP3*, *GGCT2*;*1*, *GGCT2*;*2*, and *GGCT2*;*3* were analyzed by quantitative RT-PCR using the absolute quantification method and normalized by those of *β-tubulin*. (a) Comparison of young and old leaves from the vegetative-stage samples. (b-d) Results of the reproductive stage samples. (b) Results of flowers/buds, siliques, and stems. (c) Comparison of the rosette and cauline leaves of reproductive stage samples. (d) Comparison of the upper, middle, and lower roots of the reproductive stage samples. The values and error bars represent the mean and standard deviation of four or five biological replicates. In (a) and (c), asterisks indicate significant differences (Student’s *t*-test, *P* < 0.05). In (d), different letters indicate significant differences (Tukey’s test, *P* < 0.05).

There was also a notable trait that the amount of *GGP3* transcript was significantly higher in mature leaves than in young leaves at the vegetative stage (Fig. **2a**), and a similar tendency was observed for *GGP1* (*P* < 0.1). As mentioned above, the *GGP1* transcript was highly abundant when the leaves achieved a rosette arrangement at the reproductive stage (Fig. **2b,c**). In contrast, the amount of *GGCT2;2* transcript was significantly higher in young leaves than mature ones at the vegetative stage (Fig. **2a**).

A similar trait was confirmed for the reproductive stage wherein the *GGCT2*;*2* transcript in cauline leaves was significantly higher than that in rosette leaves (Fig. **2c**). Furthermore, the *GGCT2*;*2* transcript in the lower roots, which is considered as the youngest in roots, was significantly larger than in the middle and upper roots (Fig. **2d**).

### Changes in expression of *GGP* and *GGCT* genes under sulfur and nitrogen deficiency

We also investigated the expression of *GGP1*, *GGP3*, *GGCT2*;*1*, *GGCT2*;*2* and *GGCT2*;*3* under sulfur or nitrogen deficiency because GSH functions in the storage of organic sulfur and nitrogen, and in fact, the expression of *GGCT2*;*1* sharply increases under sulfur deficiency (Joshi *et al*., 2019). The changes in expression of *GGP1*, *GGP3*, *GGCT2*;*1*, *GGCT2*;*2* and *GGCT2*;*3* in wild-type plants under nitrogen or sulfur deficiency are shown in Fig. **3**. The mRNA level of *GGCT2*;*1* was drastically increased under sulfur deficiency by 31-fold compared with that under the control condition and that of *GGCT2*;*2* was also significantly increased by 2.3-fold.

**Fig. 3.**
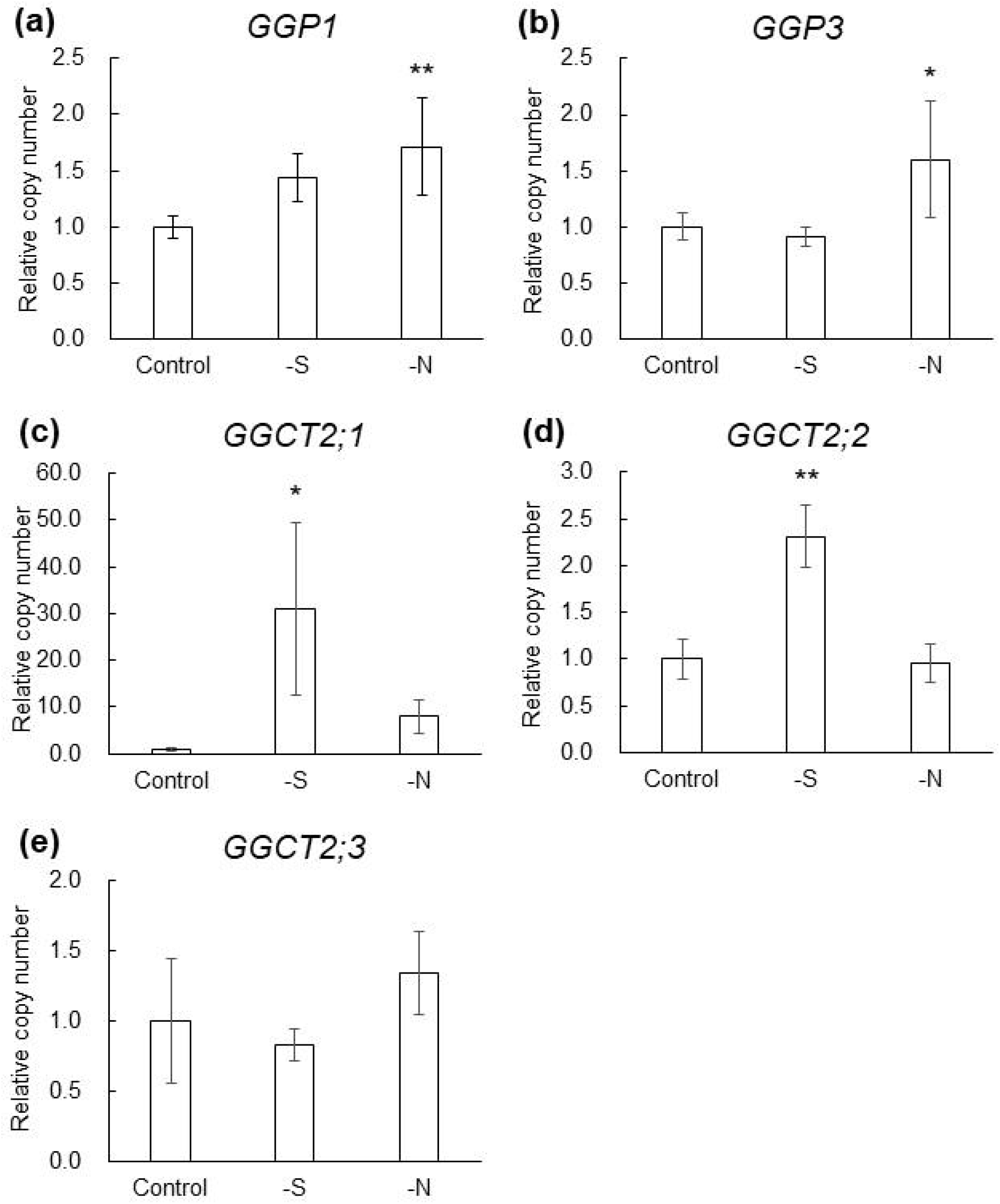
Changes in expression of the genes *GGP1*, *GGP3*, *GGCT2*;*1*, *GGCT2*;*2*, *GGCT2*;*3* under sulfur and nitrogen deficiency in wild-type plants. Plants were grown in flask culture. Expression levels under various nutritional conditions were measured by quantitative RT-PCR using the absolute quantification method, normalized by those of *β-tubulin*, and shown as control at 1. The values and error bars represent the mean and standard deviation of four to six biological replicates. Asterisks indicate significant differences from the control (Dunnett’s test, \**P* < 0.05, \*\**P* < 0.01). The expression levels of (a) *GGP1*, (b) *GGP3*, (c) *GGCT2*;*1*, (d) *GGCT2*;*2*, and (e) *GGCT2;3* are shown.

### GSH accumulation in the *ggp1* and *ggct2*;*1* knockout mutants

To confirm the contribution of GGP1 and GGCT2;1 to GSH degradation *in planta*, GSH concentration in the *ggp1-1* knockout mutant (GK-319F10, Geu-Flores *et al*., 2011), *ggct2*;*1-2* knockout mutant (SALK_56007, Joshi *et al*., 2019), and their corresponding wild-type plants (Columbia-0) was measured using 14 d-old liquid cultured seedlings. Under normal conditions, GSH concentration in *ggp1-1* was significantly higher than that in the wild-type plants, whereas *ggct2*;*1-2* did not show any significant difference (Fig. **4**), suggesting that GGP1 degrades GSH under normal conditions.

**Fig. 4.**
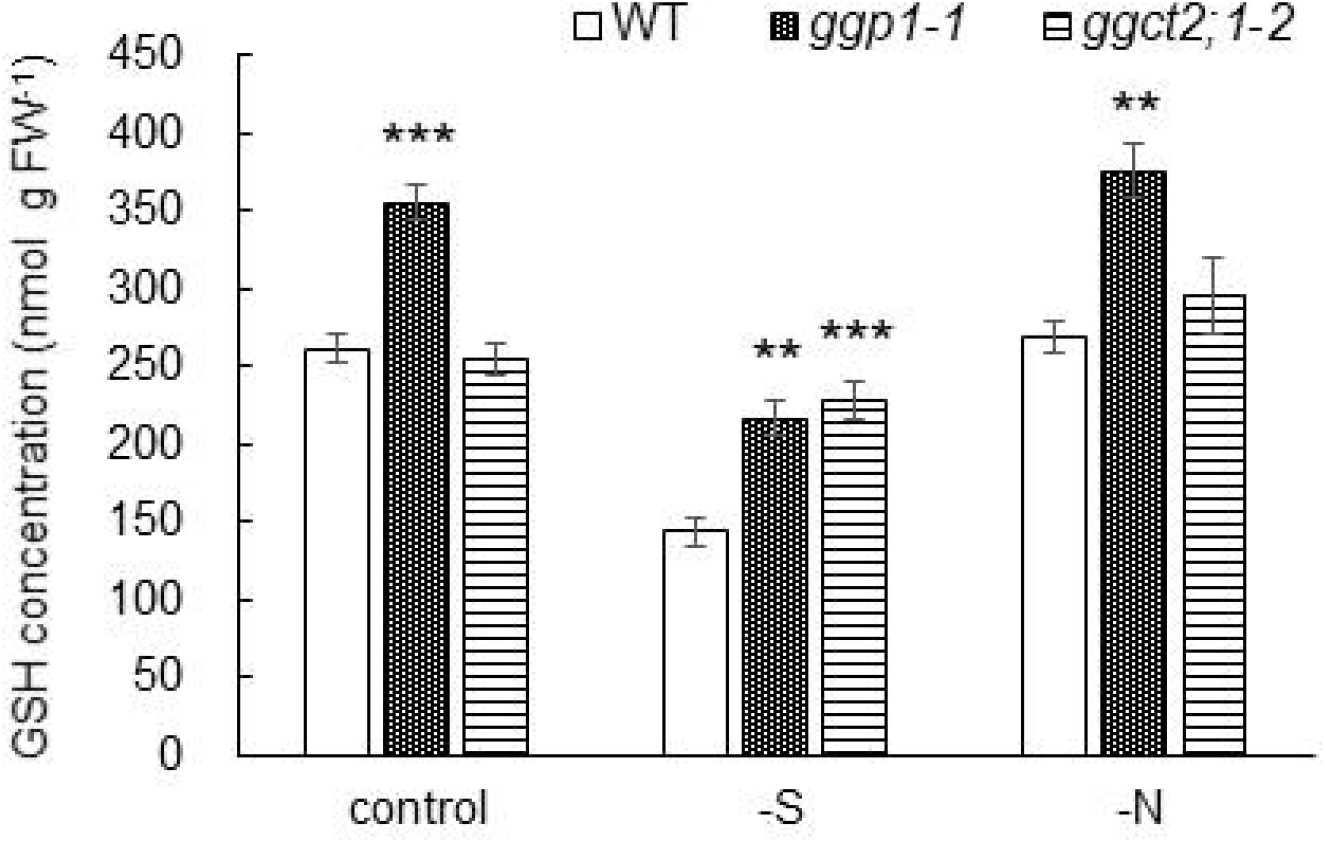
GSH concentrations in wild-type (WT) plants and *ggp1-1* and *ggct2*;*1-2* mutant plants grown for 14 d in liquid culture. The GSH concentrations in the seedlings are presented. The values and error bars represent the mean and standard deviation of four biological replicates. Asterisks indicate significant differences from the wild-type (Dunnett’s test, \**P* < 0.05, \*\**P* < 0.01).

We also measured GSH concentrations in mutants under sulfur- or nitrogen-deficient conditions. Under sulfur deficiency, the *ggp1-1* and *ggct2*;*1-2* mutants showed significantly higher GSH levels than the wild-type plants (Fig. **4**). Higher GSH levels under sulfur deficiency have also been reported for *ggct2*;*1-1* (SALK_117578), another T-DNA insertion mutant of *GGCT2*;*1* (Joshi *et al*., 2019). However, under nitrogen deficiency, the elevation of GSH levels was only observed in *ggp1-1* and not in *ggct2*;*1-2*, which was similar to normal conditions. GSH accumulation by the deletion of *GGP1* was also confirmed for *ggp1-2* (SALK_089634), another T-DNA insertion mutant of *GGP1*, under normal and sulfur- and nitrogen-deficient conditions (Fig. **S5**). These results suggest that GGP1 constantly degrades GSH, and GGCT2;1 also breaks it down under sulfur deficiency.

### mRNA levels of *GGP* and *GGCT* genes in wild-type, *ggp1-1*, and *ggct2*;*1-2* mutant plants under the control condition, sulfur or nitrogen deficiency

Homologous proteins often complement the lack of protein function. Therefore, the expression of other *GGP* and *GGCT* genes in *ggp1-1* and *ggct2*;*1-2*, as well as in wild-type plants, was investigated to determine whether this is the case for GGP1 or GGCT2;1, (Fig. **5**). The mRNA level of *GGP3* increased in the *ggp1-1* mutant plants compared with that in wild-type plants under sulfur deficiency (Fig. **5b**), suggesting a possibility that GGP3 functionally complements GGP1 under these conditions. The mRNA level of *GGCT2*;*2* increased in *ggct2*;*1-2* mutant plants compared with that in wild-type plants under sulfur deficiency, suggesting a possibility that GGCT2;2 functionally complements GGCT2;1 under these conditions.

**Fig. 5.**
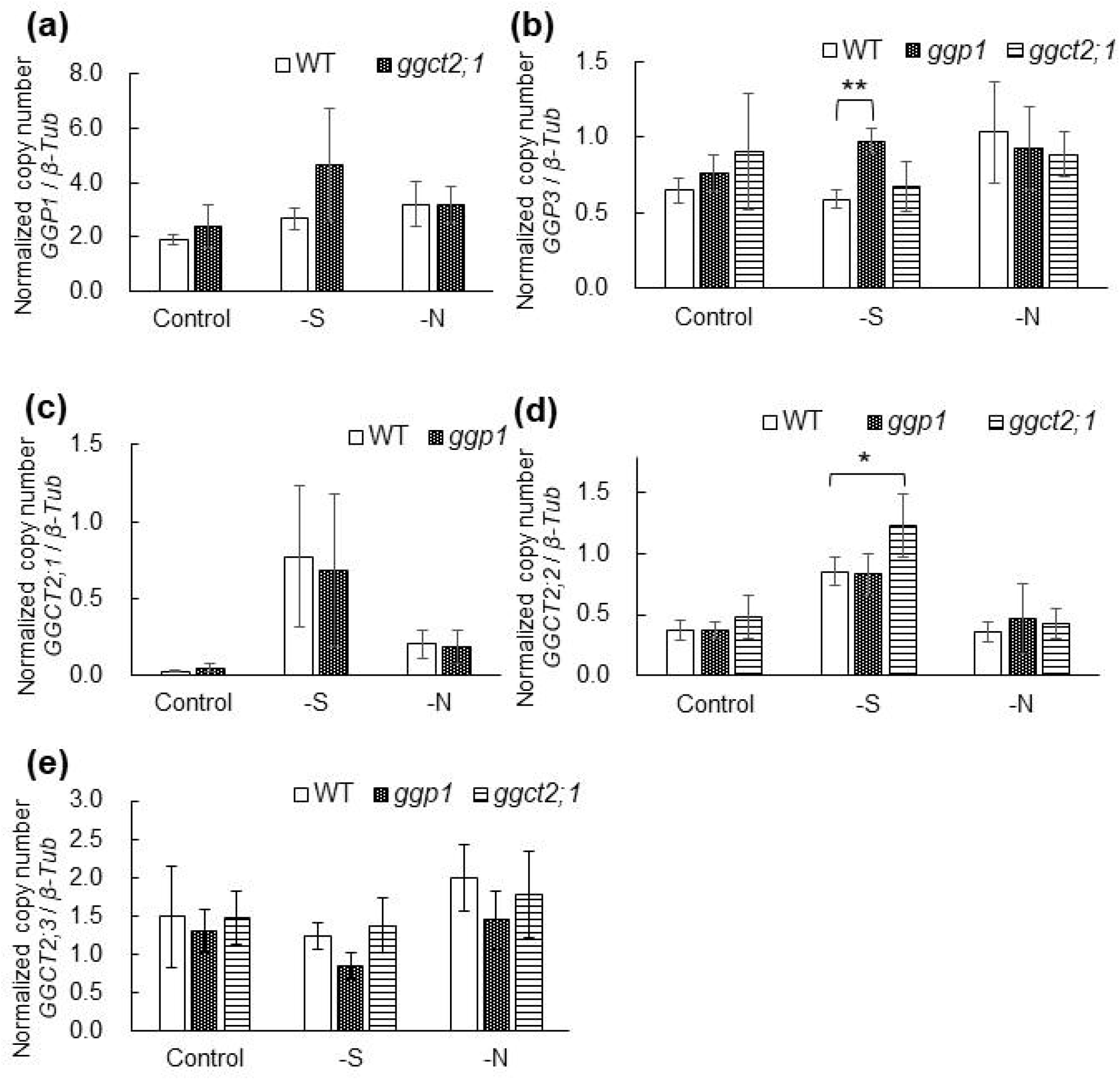
Normalized copy numbers of *GGP* and *GGCT* transcripts in wild-type (WT) plants and *ggp1-1* and *ggct2*;*1-2* mutant plants under control condition and sulfur and nitrogen deficiency. The plants were grown in liquid culture for 14 d. The entire body of the 14 d-old plants was sampled. Transcript copy numbers of *GGP1*, *GGP3*, *GGCT2*;*1*, *GGCT2*;*2*, and *GGCT2*;*3* were analyzed by quantitative RT-PCR using the absolute quantification method and normalized by those of *β-tubulin*. The values and error bars represent the mean and standard deviation of four to six biological replicates. Asterisks indicate significant differences compared to WT (Dunnett’s test, \**P* < 0.05, \*\**P* < 0.01). Relative copy numbers of (a) *GGP1*, (b) *GGP3*, (c) *GGCT2*;*1*, (d) *GGCT2*;*2*, and (e) *GGCT2*;*3* are shown.

## Discussion

### Newly discovered GSH degradation activity of GGP1 and GGP3

In the present study, we found that GGPs, enzymes that were originally reported to process GSH-conjugates (Geu-Flores *et al*., 2009), also degrade GSH. The yeast mutants defective in GSH degradation—*dug2Δ* or *dug3Δ*—were complemented by *Arabidopsis GGP1* (Fig. **1**) and grew on the medium where GSH was the sole sulfur source. In addition, the recombinant GGP1 and GGP3 proteins showed activity for GSH degradation (Fig. **S3**; Table **1**). The *Km* values of GGP1 and GGP3 for GSH were 5.0 mM and 3.7 mM, which are reasonable because cytosolic GSH concentration ranges from 2.8 mM to 4.5 mM (Koffer *et al*., 2013). This reasonability is also confirmed by the fact that the *K_m_* values of GGCTs are 1.9 mM for GGCT2;1 (Paulose *et al*., 2013), 1.7 mM for GGCT2;2 (Kumar *et al*., 2015), and 6.7 mM (Table **1**) or 4.9 mM (Kumar *et al*., 2015) for GGCT2;3. The activity of GGP1 and GGP3 was moderate because their *k_cat_* values were 1.1-fold and 3.8-fold higher than that of GGCT2;3 (Table **1**), respectively, whereas the *k_cat_* value of GGCT2;2 is 5.7-fold higher than that of GGCT2;3 (Kumar *et al*., 2015).

Another remarkable trait of GGPs is the abundance of *GGP1* transcripts. The *GGP1* transcript was the largest among *GGPs* and *GGCTs* in the reproductive stage rosette leaves (Fig. **2**), which occupied the highest volume in the plant body. The *GGP1* transcript was also the most abundant in 2-week-old liquid cultured plants (Fig. **5**). Additionally, in the TAIR database (http://www.arabidopsis.org/), the number of expressed sequence tags (ESTs) associated with GGP1, GGP3, GGCT2;1, GGCT2;2 and GGCT2;3 were 335, 66, 13, 118, and 52, respectively, and five clones that complemented the *dug2Δ* or *dug3Δ* mutants in our screening all corresponded to *GGP1*.

Considering these facts, it is highly possible that GGPs, especially GGP1, substantially contribute to GSH degradation *in planta*. The *ggp1* knockout mutant significantly accumulated GSH compared with the wild-type plants (Fig. **4**). Therefore, GGP1 and GGP3, in addition to GGCT2;1, GGCT2;2, and GGCT2;3, should be considered when examining GSH degradation in the cytosol. In plants, the turnover rate of GSH is so fast that 80% of GSH is degraded in 1 d, and a large portion of this degradation is estimated to be cytosolic (Ohkama-Ohtsu *et al*., 2008). It is a future challenge to investigate whether GGPs and GGCTs can explain this high degradation rate.

It should also be noted that GSH degradation by GGPs is reasonable in terms of energy efficiency. This is because the pathway through GGPs does not require ATP for Glu production from GSH (Geu-Flores *et al*., 2011), whereas the pathway through GGCTs requires ATP to convert 5-oxoproline to Glu (Ohkama-Ohtsu *et al*., 2008). Thus, it is considered reasonable for plants to use GGPs when degrading GSH.

### Bifunctionality of GGP1 and GGP3 in GSH degradation and glucosinolate synthesis

GGP1 and GGP3 were initially reported as enzymes that process GSH-conjugates in the biosynthesis of glucosinolates and camalexin (Geu-Flores *et al*., 2009); therefore, the competition between the reactions can be considered. The *K_m_* value of GGP1 was 5.0 mM for GSH (Table **1**), which is appropriate considering the cytosolic GSH concentration mentioned above. In contrast, the *K_m_* value for a GSH-conjugate *S*-[(*Z*)-phenylaceto-hydroximoyl]-L-glutathione (GS-B) is 37 μM, which also suggests that GS-B can be a physiological substrate (Geu-Flores *et al*., 2009). Therefore, both reactions are considered to occur in plants, competing with each other. The ATTED-II database (Obayashi *et al*., 2018, https://atted.jp/) shows that *GGP1* is coregulated with genes involved in glucosinolate synthesis pathways and the sulfur assimilation pathway, such as *APS1* (*ATP sulfurylase 1*, *At3g22890*), *APR1* (*APS reductase 1*, *AT4G04610*), *APK1* (*APS kinase 1*, *AT2G14750*), *APK2* (*APS kinase 2*, *At4g39940*), *APS3* (*ATP sulfurylase 3*, *AT4G14680*), and *SERAT2*;*2* (*serine acetyltransferase2*;*2*, *At3g13110*). Because GSH is the primary storage form of organic sulfur in cells, it is conceivable that GSH degradation, together with sulfur assimilation, supplies organic sulfur when the demand is high.

If GGPs process both GSH and GSH-conjugates, the priority of the reactions likely varies depending on the substrate concentration and is considered reasonable in plants. The affinity of GGPs for GSH-conjugates is much higher than for GSH; when GSH-conjugates accumulate, it is likely that processing GSH-conjugates will be prioritized over GSH degradation, leading to an automatic reduction in GSH degradation. Although it is well known that glucosinolates and camalexin function in alleviating biotic stresses, high concentrations of GSH are also crucial for resistance to viruses, bacteria, and fungi because GSH functions in ROS scavenging and signaling (Hernández *et al*., 2017). Therefore, it is very reasonable for plants experiencing biotic stress to synthesize glucosinolates and camalexin and simultaneously decrease GSH degradation. Additionally, it is notable that *Ralstonia solanacearum*, a pathogenic bacterium, produces the virulence effector protein RipAY, which has GGCT activity (Mukaihara *et al*., 2016). GSH degradation by RipAY causes the depletion of the GSH pool in plant cells, increasing susceptibility in the early stages of infection.

In contrast, GSH degradation may be prioritized over GSH-conjugate degradation under sulfur starvation because the transcript levels of upstream genes in glucosinolate synthesis pathways are downregulated under sulfur-deficient conditions (Aarabi *et al*., 2016), and thus, the concentrations of GSH-conjugates are presumed to decrease. It is reasonable that GSH degradation activates under sulfur deficiency because GSH degradation functions as the distribution of stored organic sulfur. In addition, it has recently been shown that glucosinolates are degraded under sulfur deficiency to reallocate sulfur atoms in them, and that GSH is required for the glucosinolate degradation pathway (Sugiyama *et al*., 2021). Therefore, glucosinolate synthesis by GGPs, which consumes GSH, may be inactivated under sulfur deficiency, and GSH is possibly utilized for glucosinolate degradation. Overall, bifunctionality of GGPs potentially contribute to sulfur flow shift under sulfur deficiency from sulfur consumption in glucosinolate synthesis to sulfur reallocation through GSH and glucosinolate degradation.

Incidentally, *GGP3* is not coregulated with genes involved in glucosinolate synthesis pathways or the sulfur assimilation pathway. This is possibly because *GGP3* is a redundant gene of *GGP1* and its function is limited, considering the expression of *GGP3* was slightly but significantly promoted in the *ggp1* knockout mutant under sulfur-deficient conditions (Fig. **5b**), where there was high demand for Cys. GGP1 is presumed to contribute to GSH degradation more than GGP3 because *GGP1* tends to be expressed more actively than *GGP3* (Fig. **2**), corresponding with the fact that 335 ESTs are associated with *GGP1*, whereas *GGP3* has only 66 ESTs according to TAIR.

### Expression of *GGP1* is active in mature organs and that of *GGCT2*;*2* is active in young organs

The results of the complementation assay of yeast and *in vitro* activity assay showed that GGP1, GGP3, GGCT2;1, GGCT 2;2, and GGCT 2;3 solely degraded GSH without forming heterodimers (Fig. **1**; Table **1**). In addition, GGCT2;1 (Paulose *et al*., 2013) are localized in the cytosol and GGCT 2;2 and GGCT 2;3 are also predicted to be cytosolic in the TAIR database, so it is interesting to consider their functional differences.

The expression patterns of *GGPs* and *GGCTs* differed depending on the organ type (Fig. **2**). The expression of *GGP1* and *GGP3* was more active in mature organs than in young ones, like that in mature leaves than young leaves (Fig. **2a**), and in rosette leaves than cauline leaves (Fig. **2c**). More specifically, GGP1 is localized in the vascular tissues of leaves (Geu-Flores *et al*., 2011). Therefore, GSH degradation by GGP1 is considered functional in the distribution of constituent amino acids in the vascular tissue of leaves, contributing to the subsequent synthesis of proteins and other metabolites. For instance, Cys is the precursor molecule of numerous sulfur-containing metabolites such as methionine, vitamins, cofactors, and Fe-S clusters. Methionine can also be processed into *S*-adenosyl-L-methionine or ethylene (Romero *et al*., 2014). Therefore, the contribution of GGP1 to the synthesis of proteins and their metabolites should be further studied to fully understand the physiological significance of GGP1.

In contrast, the expression of *GGCT2*;*2* was more active in young organs than in old ones, such as in young leaves than mature leaves (Fig. **2a**), in cauline leaves than rosette leaves (Fig. **2c**), and in lower roots than middle or upper roots (Fig. **2d**). These data mostly coincide with public microarray data visualized by Genevestigator (https://www.genevestigator.com/) (Fig. **S6**). GGCT2;2 has a high affinity and high activity compared with other GGPs and GGCTs. The *K_m_* of GGCT2;2 is 1.7 mM (Kumar *et al*., 2015), which is lower than that of GGP1 (5.0 mM, Table **1**), GGP3 (3.7 mM, Table **1**), and GGCT2;3 (6.7 mM, Table **1**; 4.9 mM, Kumar *et al*., 2015). The *k_cat_* of GGCT2;2 is *c*. 5.7-fold higher than that of GGCT2;3 (Kumar *et al*., 2015), whereas the *k_cat_* of GGP1 was almost the same as that of GGCT2;3, and the *k_cat_* of GGP3 was 3.8-fold higher than GGCT2;3 (Table **1**). Therefore, GSH in young organs is presumed to be actively degraded by GGCT2;2. The physiological function of GGCT2;2 is possibly to provide developing organs with constituent amino acids to support plant growth. The involvement of GGCT2;2 in plant growth can be inferred from the fact that *GGCT2;1—*a gene explicitly expressed under sulfur deficiency—is related to root architecture under sulfur deficiency (Joshi *et al*., 2019). It is interesting to investigate whether GGCT2;2 is related to root architecture under normal conditions.

### The model of cytosolic GSH degradation in plants under normal and sulfur-deficient conditions

Our results of *in vitro* activity assays of GGP1, GGP3, and GGCT2;3 (Fig. **S3**; Table **1**) together with the previous reports on GGCT2;1 (Paulose *et al*., 2013), GGCT2;2, and GGCT2;3 (Kumar *et al*., 2015) show that all of these enzymes can be functional under physiological GSH concentrations in the cytosol of plants. Therefore, we developed a model of cytosolic GSH degradation (Fig. **6**), mainly based on transcript levels.

**Fig. 6.**
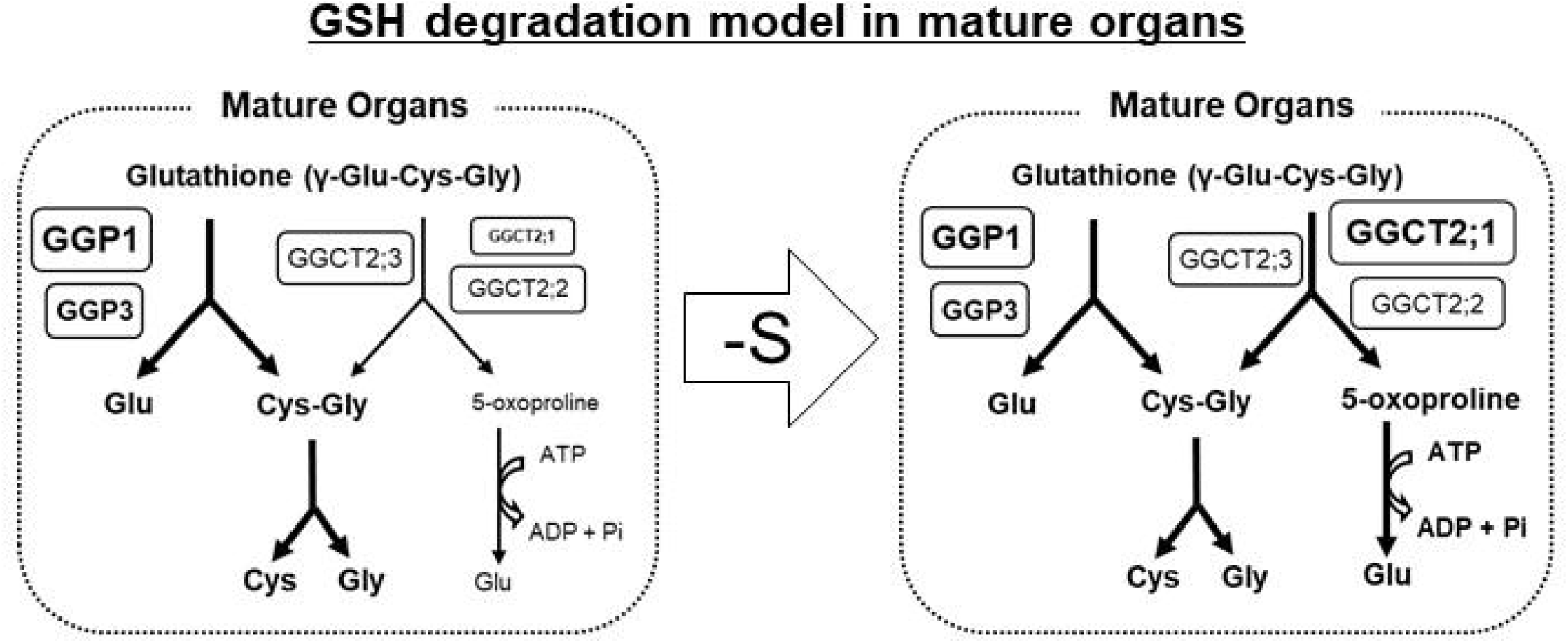
Model of cytosolic degradation of GSH in plants. GGP1 is considered to be the main enzyme which degrades GSH in mature organs, and GGP3 is presumed to support it. GGCT2;1 accelerates the degradation under sulfur deficiency. The expression of GGCT2;3 is constitutive.

The GGP1 transcript was the most abundant in mature organs among GGPs and GGCTs (Fig. **2**, **5**), which is consistent with the high number of GGP1 ESTs in the TAIR database and the fact that all clones that complemented the *dug2Δ* and *dug3Δ* yeast mutants corresponded to *GGP1*. In our mutant analysis, *ggp1* knockout mutants showed GSH accumulation under normal and sulfur- and nitrogen-deficient conditions (Fig. **4**, **S5**). Therefore, we assumed that GGP1 generally contributes to GSH degradation in mature organs (Fig. **6**). Under sulfur deficiency, the transcript levels of *GGCT2*;*1* and *GGCT2*;*2* increased by 31-fold and 2.3-fold, respectively (Fig. **3c**,**d**), so their contribution, especially that of GGCT2;1, is presumed to increase (Fig. **6**). Because GSH concentration is low under sulfur deficiency, GGCT2;1 and GGCT2;2, which have a higher affinity for GSH than GGP1, GGP3, and GGCT2;3 (Table **1**; Paulose *et al*., 2013; Kumar *et al*., 2015), may be more efficient under these conditions. In addition, *GGCT2*;*2* transcripts were abundant under normal conditions in developing organs such as young leaves and root tips (Fig. **2**, **S7**); therefore, the contribution of GGCT2;2 is considered larger in these organs.

The degradation pathway through GGPs is more efficient than GGCTs because the former directly produces Glu without ATP (Geu-Flores *et al*., 2011), whereas the latter requires ATP to convert 5-oxoproline to Glu (Ohkama-Ohtsu *et al*., 2008). Thus, it is conceivable that plants fundamentally degrade GSH with GGPs and utilize GGCTs only when the demand for GSH degradation is high; thus, high-affinity or high-activity enzymes are needed in conditions where organs are developing or under sulfur deficiency. Notably, our findings suggest that plants utilize two types of GSH degradation pathways to meet the demand for constituent amino acids with less energy.

## Acknowledgements

We thank Dr. Akira Nozawa (Ehime University) and Dr. Toru Fujiwara (The University of Tokyo) for kindly providing pFL61 and the method for yeast transformation. This work was partly supported by JSPS KAKENHI Grant Number 15KT0028, 16K07639 and 19H02859 to N.O-O.

## Author contributions

T.I., T.K., K.N., M.U. and N. O-O. designed the experiments and analyzed the data; T.I., T.K., K.N., M.U., S.M., S.A. and K.K. performed experiments; A. M-N., R. S., M.Y.H., T.Y. and N.O-O. supervised the experiments; T.I., A. M-N., R. S., M.Y.H. and N.O-O. contributed to the writing of the manuscript.

## Data availability

The data that supports the findings of this study are available in the supplementary material of this article.

