## Supplementary data for "Characterization of γ-Glutamyl Peptidases and γ-Glutamyl Cyclotransferases for Glutathione Degradation in *Arabidopsis*"

**Table S1** List of yeast strains used in this study.

| Strain | Description | Genotype |
|---|---|---|
| Y00000 | BY4741 | *MAT a; his3Δ1; leu2Δ0; met15Δ0; ura3Δ0::kanMX4* |
| Y05729 | *dug2Δ* | *BY4741; MAT a; his3Δ1; leu2Δ0; met15Δ0; ura3Δ0; YBR281c::kanMX4* |
| Y02021 | *dug3Δ* | *BY4741; MAT a; his3Δ1; leu2Δ0; met15Δ0; ura3Δ0; YNL191w::kanMX4* |

**Table S2** Primers used for constructing and sequencing expression plasmids for yeast.

| Primers | Sequences (5' to 3') |
|---|---|
| pGK5 | TACAGATCATCAAGGAAGTAATTATC |
| pGK3 | TTAGCGTAAAGGATGGG |
| *Eco*RI-At4g30530-F | TATGGCGAATTCATGGTGGAGCAAAAGAGATACG |
| *Xho*I-At4g30530-R | ATTGTACT GAGCTCTAGTTAGTTGGAACTCTGCC |
| *Bam*HI-At1g44790-F | CGGGATCCCGATGGCGATGTGGGTATTCGGGT |
| *Eco*RI-At1g44790-R | TATGGCGAATTC CTAGACATTGTTGGCGGTGGC |
| *Bam*HI-At4g31290-F | CGGGATCCCGATGGTGATGTGGGTCTTTGGCT |
| *Eco*RI-At4g31290-R | TATGGCGAATTCTTATATTGTAGTGGCAACAGCTTCTGG |
| *Bam*HI-At5g26220-F | CGGGATCCCGATGGTTTTGTGGGTATTTGGATATGGTTC |
| *Eco*RI At5g26220-R | TATGGCGAATTCTCATGATGCAAAGACCCGTTGACG |

**Table S3** Primers used for the construction of *E. coli* expression strains.

| Gene | Forward primers (5' to 3') | Reverse primers (5' to 3') |
|---|---|---|
| *GGP1* | CACCATGGTGGAGCAAAAGAGATA | CTAGTTAGTTGGAACTCTGCCT |
| *GGP3* | CACCATGGTGGTTATTGAGCAGAA | TCAACCTTTCAGGAAGTTTTTG |
| *GGCT2;1* | CACCATGGTTTTGTGGGTATTTGGATATGG | TCATGATGCAAAGACCCGTTGACGA |
| *GGCT2;2* | CACCATGGTGATGTGGGTCTTTGGC | TTATATTGTAGTGGCAACAGC |
| *GGCT2;3* | CACCATGGCGATGTGGGTATTCGG | CTAGACATTGTTGGCGGTGGCGG |
| *M13* | GTAAAACGACGGCCAG | CAGGAAACAGCTATGAC |

**Table S4** Primers used for the quantitative PCR.

| Gene | Forward primers (5' to 3') | Reverse primers (5' to 3') |
|---|---|---|
| *GGP1* | CTACGTCAAGCAAGAATTTGCT | GCCAAATAGATAATTTGGGAGTG |
| *GGP3* | CGGCCACCAGATAATTACTAGA | ATGCTGGGACCTCAACGT |
| *GGCT2;1* | GGTTTTGATTTTGATGAAAAACTC | CACCCCAGCAAATAGCTC |
| *GGCT2;2* | GGATTTCACTACGATGAGAAAGTG | TACCCCAGCATATGGCTT |
| *GGCT2;3* | GCCAAACAGATAGTTAAAGCG | CGGATTCTGAAAGAATGTGC |
| *TUB4* | GCTCGCTAATCCTACCTTTGG | AGCCTTGGGAATGGGATAAG |

**Table S5** Primers used for screening *ggp1* and *ggct 2;2* mutant and mRNA analysis.

| Primers | Sequences (5' to 3') |
|---|---|
| GK-319-F | GTAAATCACACTTCTTGGTTTGG |
| GK-319-R | CTTTGTTATACTCAGGATGTCCTTG |
| GABI-8474 | ATAATAACGCTGCGGACATCTACATTTT |
| At4g30530F | GGCTACGTCAAGCAAGAATT |
| At4g30530R | CAAATAGATAATTTGGGAGTGAAA |
| SALK_089634F | GCCTTGCTGGTATAAACTTATG |
| SALK_089634R | TGACAACTGCAATTATGGCT |
| GGCT2;1F | CAATCACATACTCTTCCTCATGC |
| GGCT2;1R | GGAACGAAGAGAGTGTGAATAC |
| pROKr3 | CCTTTCGCTTTCTTCCCTTCCTTTCT |
| At5g26220F | CAATCACATACTCTTCCTCATGC |
| At5g26220R | GGAACGAAGAGAGTGTGAATAC |

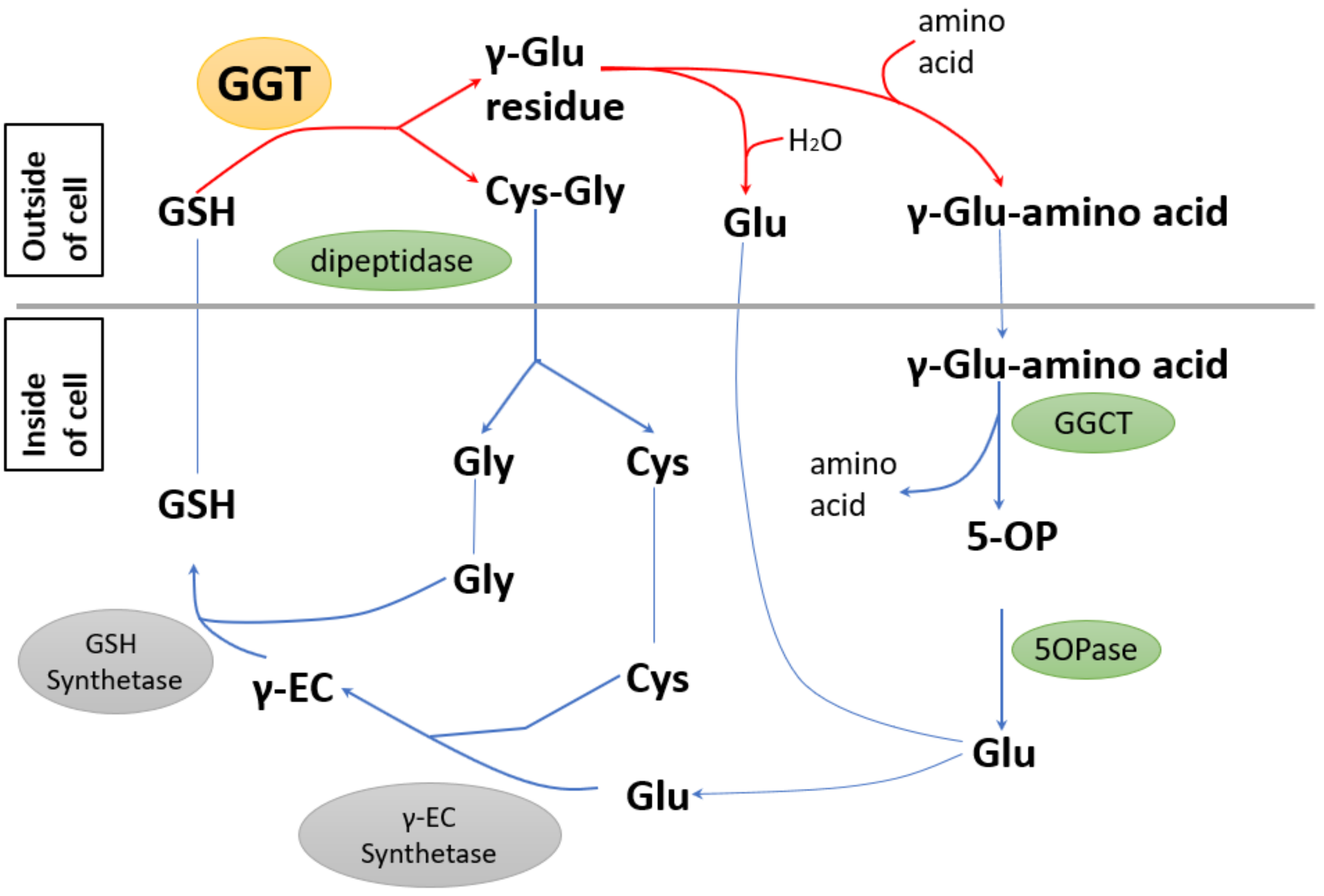


**Fig. S1** The γ-glutamyl cycle proposed in mammals. GGT; γ-glutamyl transpeptidase, GGCT; γ-glutamyl cyclotransferase, 5OPase; 5-oxoprolinase, GSH; glutathione, γ-EC; γ-glutamyl cysteine, 5OP; 5-oxoproine

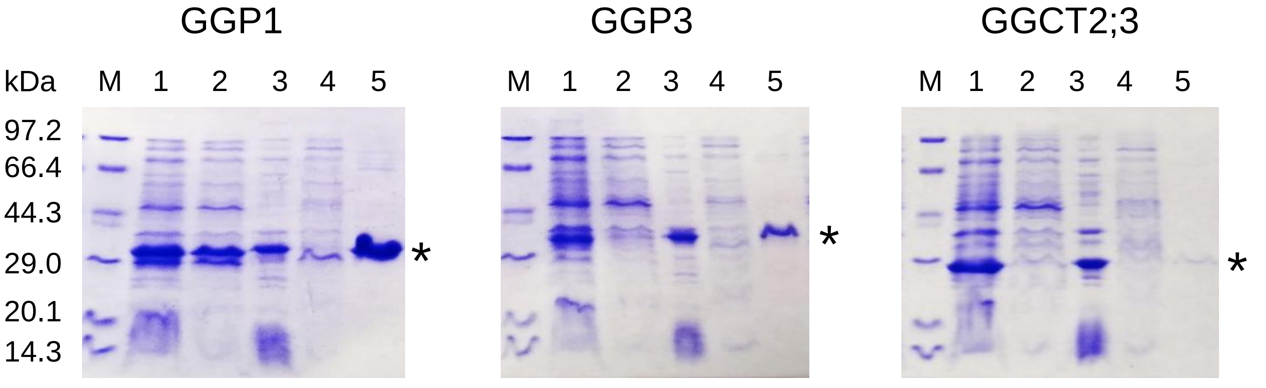


**Fig. S2** Confirmation of the result of protein purification by SDS-PAGE. N-terminal His tagged proteins were expressed in *E.coli* and purified with nickel columns. M: marker; lane 1: crude protein extract; lane 2: soluble fraction; lane 3: insoluble fraction; lane 4: flow-through; lane 5: purified protein. Asterisks indicate the size of the purified proteins.

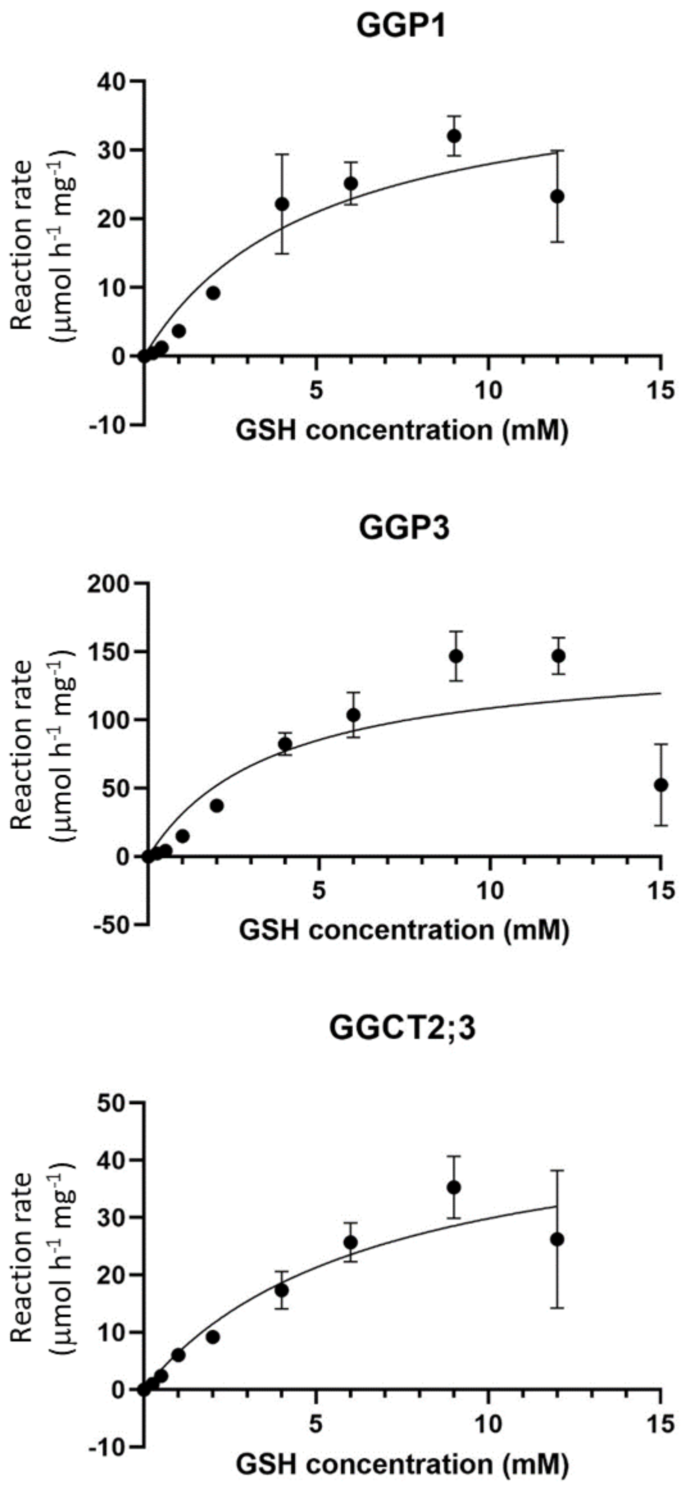


**Fig. S3** Michaelis-Menten plot of GSH degradation activity of GGP1, GGP3, and GGCT2;3. Recombinant GGP1, GGP3, and GGCT2;3 were incubated in 50 µl of reaction mixture containing 0.25 mM to 15.0 mM GSH for 30 min at 37 °C. The points and error bars represent the means and standard deviations of three biological replicates, and kinetic parameters were calculated using GraphPad Prism 9.

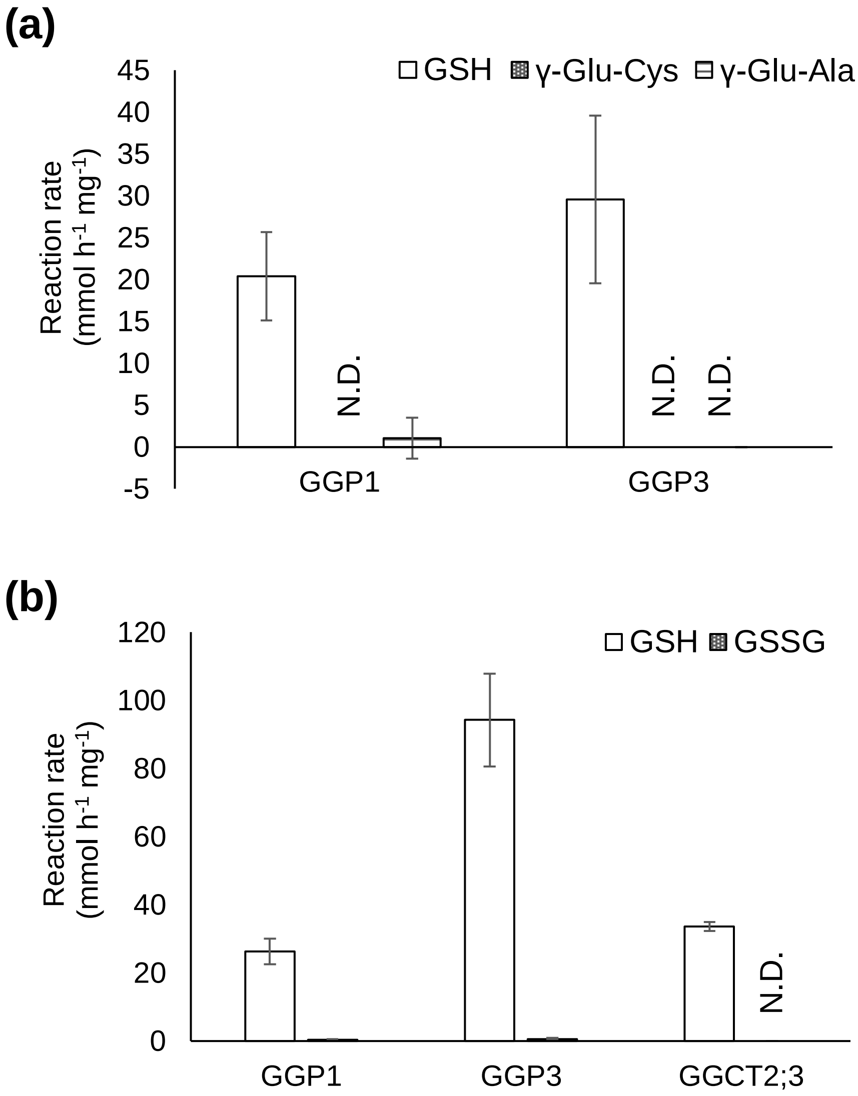


**Fig. S4** Degradation activity towards other γ-glutamyl compounds. Recombinant GGP1, GGP3, and GGCT2;3 were incubated in 50 μl of reaction mixture containing various γ-glutamyl compounds for 30 min at 37 °C. (a) Degradation activity for GSH, γ-Glu-Cys, and γ-Glu-Ala. (b) Degradation activity for GSH and GSSG. The values and error bars represent the mean and standard deviation of three biological replicates.

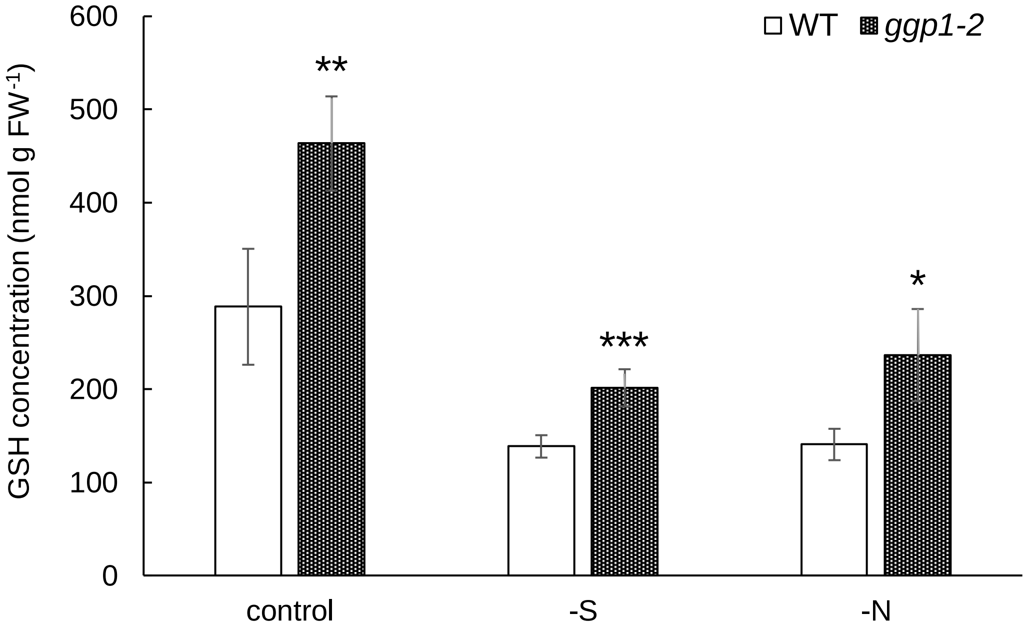


**Fig. S5** GSH concentrations in wild-type (WT) and *ggp1-2* mutant plants grown for 14 days in liquid culture. GSH concentrations in seedlings are presented. The values and error bars represent means and standard deviations of four to six biological replicates. Asterisks indicate significant differences from wild type (Student's *t*-test, $*P < 0.05$, $**P < 0.01$, $***P < 0.001$).

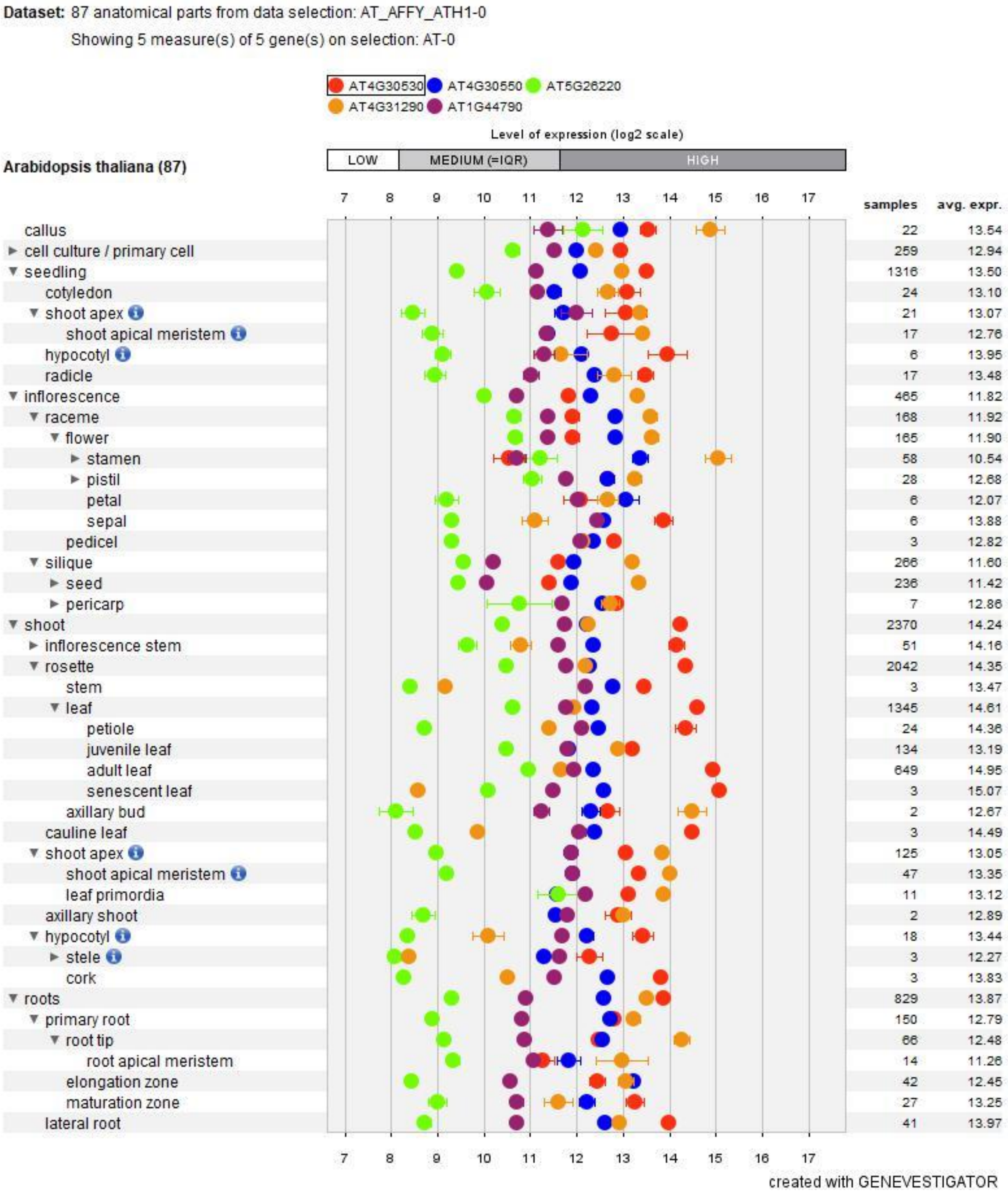


**Fig. S6** Expression analysis based on a public microarray database. The expression levels of *GGP1* (*AT4G30530*), *GGP3* (*AT4G30550*), *GGCT2;1* (*AT5G26220*), *GGCT2;2* (*AT4G31290*) and *GCT2;3* (*AT1G44790*) were visualized by Genevestigator. Error bars represent standard errors.
